# Impact of Shelterin Complex on Telomere Accessibility

**DOI:** 10.1101/2022.05.24.493289

**Authors:** Sajad Shiekh, Amanda Jack, Golam Mustafa, Sineth G. Kodikara, Prabesh Gyawali, Mohammed Enamul Hoque, Ahmet Yildiz, Hamza Balci

## Abstract

Shelterin plays critical roles in maintaining and protecting telomeres by regulating access of various physiological agents to telomeric DNA. We present single molecule measurements investigating the impact of the POT1 and a four-component shelterin complex on the accessibility of human telomeric DNA overhangs with physiologically relevant lengths (28-150 nt), which to our knowledge is the first direct approach to measure this effect on such telomeric constructs. To quantify telomere accessibility, we monitored transient binding events of a short peptide nucleic acid (PNA) probe that is complementary to telomeric overhangs using FRET-PAINT. Although POT1 has a mild G-quadruplex unfolding activity, it reduced accessibility of the PNA probe by ∼2.5 fold, indicating that POT1 effectively binds to and protects otherwise exposed telomeric sequences. In comparison, a four-component shelterin reduced the accessibility of telomeric overhangs by ∼5-fold. This enhanced protection suggests shelterin restructures the region between single and double stranded telomere, which is otherwise the most accessible part of the overhang, by a synergistic cooperation of shelterin components located on single and double stranded telomere.

## INTRODUCTION

Telomeres are 5-10 kilobase pair long repetitive DNA sequences (GGGTTA in humans) that cap the ends of eukaryotic chromosomes and serve several critical physiological roles in protection and maintenance of chromosome ends ^1,2^. Telomeres terminate with a 50-300 nucleotide (nt) long 3′ single-stranded overhang (ssTEL), which plays an active role in successive rounds of cellular replication and DNA preservation ^2–4^. To maintain overall genomic integrity, telomeres associate with a multi-protein complex called shelterin ^5,6^. In humans, shelterin consists of six distinct proteins, TRF1, TRF2, TIN2, RAP1, POT1, and TPP1 ^5–7^. TRF1 and TRF2 recognize and bind to the double stranded telomere (dsTEL) ^8,9^, while POT1 binds to ssTEL ^10,11^. RAP1 forms a complex with TRF2 to regulate its sequence specific binding affinity ^12^. TIN2 and TPP1 orchestrate the interactions with TRF2, TRF1 and POT1, and form a direct protein-mediated link between ssTEL and dsTEL ^13,14^. Deletion or dysfunction of shelterin proteins activates various DNA damage repair pathways at telomeres ^5,7^.

Many key cellular functions are regulated by conformational dynamics and structural topologies retained by DNA in respective physiological conditions. The G-rich telomeric overhang is prone to fold into an array of G-quadruplex (GQ) structures. GQs have been reported to form in many regions of human genome including promoter regions and telomeres ^15,16^ and their formation has been established *in vitro* ^17,18^ and *in vivo* ^19–21^. There is growing evidence for roles of GQs in essential cellular processes such as DNA replication, transcription and translation ^22,23^, as well as their pivotal role in telomere maintenance ^24^. These roles have made GQs potentially important therapeutic targets ^25^.

The multiple GQs that form in ssTEL present a major hurdle for telomeric DNA synthesis and impede the physical association of telomere-binding proteins that are dedicated to protecting and modulating the organization of telomeres ^26–28^. These GQs might form compact higher order structures that further reduce the accessibility of ssTEL ^29–32^. They also help to distinguish the telomeres from DNA double strand breaks and prevent activation of superfluous DNA damage response ^1,2^. Resolving these structures requires helicase unfolding activity ^33–35^ or ssDNA binding proteins ^28,36^. POT1 binding destabilizes telomeric GQs ^37^, which might potentially make the telomeric overhang more accessible for telomerase and other enzymes ^36^. However, it may also serve to protect otherwise exposed segments of ssTEL that are not folded into a GQ structure.

The compaction of ssTEL into higher order structures via tandem GQs could potentially contribute to the robust protection of telomeres in coordination with telomere associated proteins. Here, we investigated how such an interplay and coordination between telomeric GQs and shelterin might take place using single molecule FRET-PAINT (Förster Resonance Energy Transfer-Point Accumulation in Nanoscale Topography) ^38^. In this assay, transient bindings of Cy5-labeled short peptide nucleic acid (PNA) strand to accessible regions of Cy3-labeled telomeric DNA are used to characterize the accessibility of the overhang. We previously showed that we can detect binding events of Cy5-PNA to ssTEL as long as ∼168 nt, presumably because ssTEL becomes highly compact upon tandem GQ formation ^39^. Due to their much higher stability, telomeric GQs largely inhibit binding of the short Cy5-PNA strand, which is complementary to only 7-nt of ssTEL. To illustrate, the Cy5-PNA binding frequency for an overhang that contains 4 GGGTTA repeats (which can fold into a GQ) was an order of magnitude smaller than the binding frequency for an overhang that contains a single GGGTTA repeat (cannot form a GQ), even though it contains four times as many potential binding sites. Therefore, the large majority of GQs remain folded while PNA primarily binds to sites not protected within a GQ ^39^.

In this study, we investigated the accessibility of telomeric DNA constructs of physiologically relevant lengths in the absence and presence of POT1 or a 4-component shelterin that includes TRF1, TIN2, POT1 and TPP1 (will be referred to as just ‘shelterin’ for brevity). The telomeric overhangs can fold into 1-6 tandem GQs. We characterized the frequency and location (within different parts of the telomeric overhang) of the Cy5-PNA binding events before and after introducing POT1 or shelterin. Both POT1 and shelterin reduced accessibility of Cy5-PNA although shelterin provided more efficient protection and resulted in more significant changes in Cy5-PNA accessibility maps. These findings suggest coordination between the components of shelterin that are localized on dsTEL and ssTEL play a role in reducing accessibility of the telomeric overhangs ^39^.

## RESULTS

We studied the accessibility of telomeric overhangs with 4-24 GGGTTA repeats in the absence or presence of POT1 or a 4-protein shelterin complex (POT1,TPP1,TIN2 and TRF1) using a FRET-PAINT assay (Fig. 1A, B and C) ^32,39^. Purification data and sequences for DNA constructs are presented in Supplementary Fig. S1 and Table S1. The accessible G-Tracts in ssTEL, such as those not protected in a GQ, are targeted by a complementary Cy5-PNA strand. Transient binding events of Cy5-PNA to accessible G-Tracts are observed as FRET jumps as shown in Fig. 1D. The observed FRET levels are correlated with the position of the accessible G-Tracts with respect to the ssDNA/dsDNA junction, where Cy3 is located. We previously established the detection sensitivity and applicability of FRET-PAINT method by studying long telomeric sequences ^39^. In the current study, we performed similar measurements to establish applicability of the method in the presence of POT1, which extends the overhang upon binding to it. Fig. S2 shows that despite the extension of the overhang by POT1, it is still possible to detect binding of Cy5-PNA to sites furthest from the donor molecule. Another important technical issue when performing comparative measurements with POT1 and shelterin is the necessity to keep a consistent surface quality, which can best be achieved by performing the measurements within the same sample chamber. However, that requires effective removal of POT1 molecules from the sample chamber, especially those that are bound to the telomeric overhang, before shelterin is introduced. As shown in Supplementary Fig. S3, we developed a protocol that achieves this and recovers the accessibility pattern observed in DNA-only case before shelterin is introduced.

**Figure 1:**
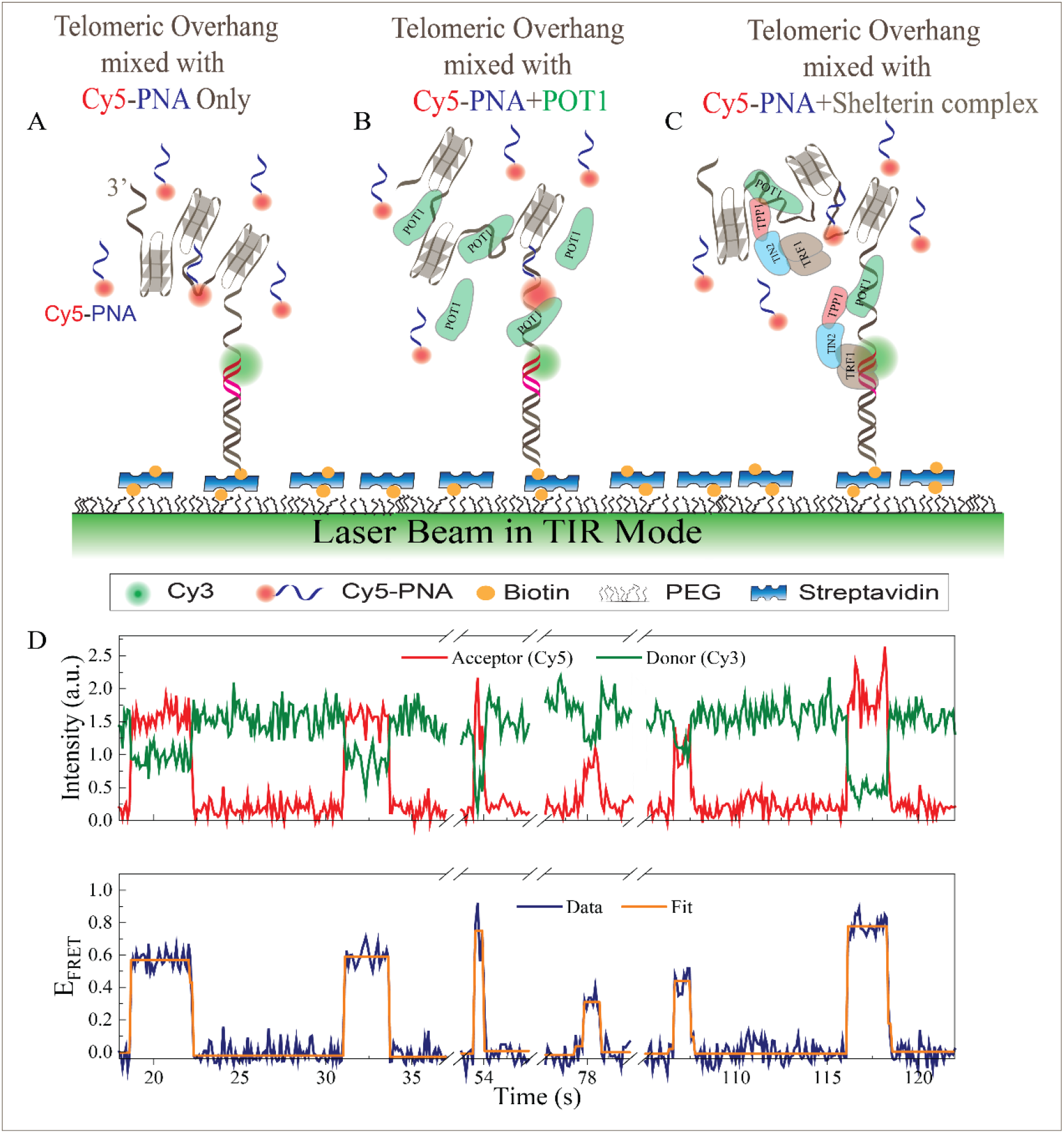
Schematic of FRET-PAINT and a representative smFRET time trace. **A**) Partial DNA duplex construct labeled with a donor fluorophore (Cy3) is immobilized on the PEGylated surface via streptavidin-biotin chemistry. The pdDNA constructs consist of dsTEL (red) and ssTEL (overhang) with multiple GGGTTA (G-Tract) repeats, which can fold into tandem GQs separated by unfolded regions of varying length. To access the unfolded regions within a telomeric overhang, a Cy5-labeled short PNA strand is used. The Cy5-PNA is complementary to a G-Tract. FRET signals with different levels are observed when Cy5-PNA transiently binds to accessible G-Tracts. Schematics of these measurements in the absence of any proteins, in the presence of POT1 and in the presence of shelterin (POT1, TPP1, TIN2 and TRF1) are shown in **(A), (B)**, and **(C)**, respectively. **D**) Representative smFRET time trace showing six Cy5-PNA binding events. The orange line is a fit to the FRET data and it is used to characterize the corresponding FRET levels.

We next monitored transient binding events of Cy5-PNA to Cy3-labeled DNA constructs with different overhang lengths in the absence or presence of 20 nM POT1 or 20 nM shelterin. Using normalized FRET distributions (also called accessibility maps), we infer how the accessible G-Tracts are distributed through the overhang. In the absence of proteins (Fig. 2A), the FRET histograms for [4n]G-Tract constructs are broadly distributed, suggesting that accessible G-Tracts are distributed throughout the overhang. The FRET distributions for [4n+2]G-Tract constructs are concentrated at high FRET levels, suggesting that most accessible sites are in the vicinity of ssDNA/dsDNA junction region. However, POT1 reduces the contrast between the accessibility maps of [4n]G-Tract and [4n+2]G-Tract constructs, and shelterin largely eliminates these differences. Fig. 2D-F show the contour plots of the FRET distributions in Fig. 2A-C, respectively, where the changes in the accessibility maps in the presence of POT1 or shelterin are clearly visible.

**Figure 2:**
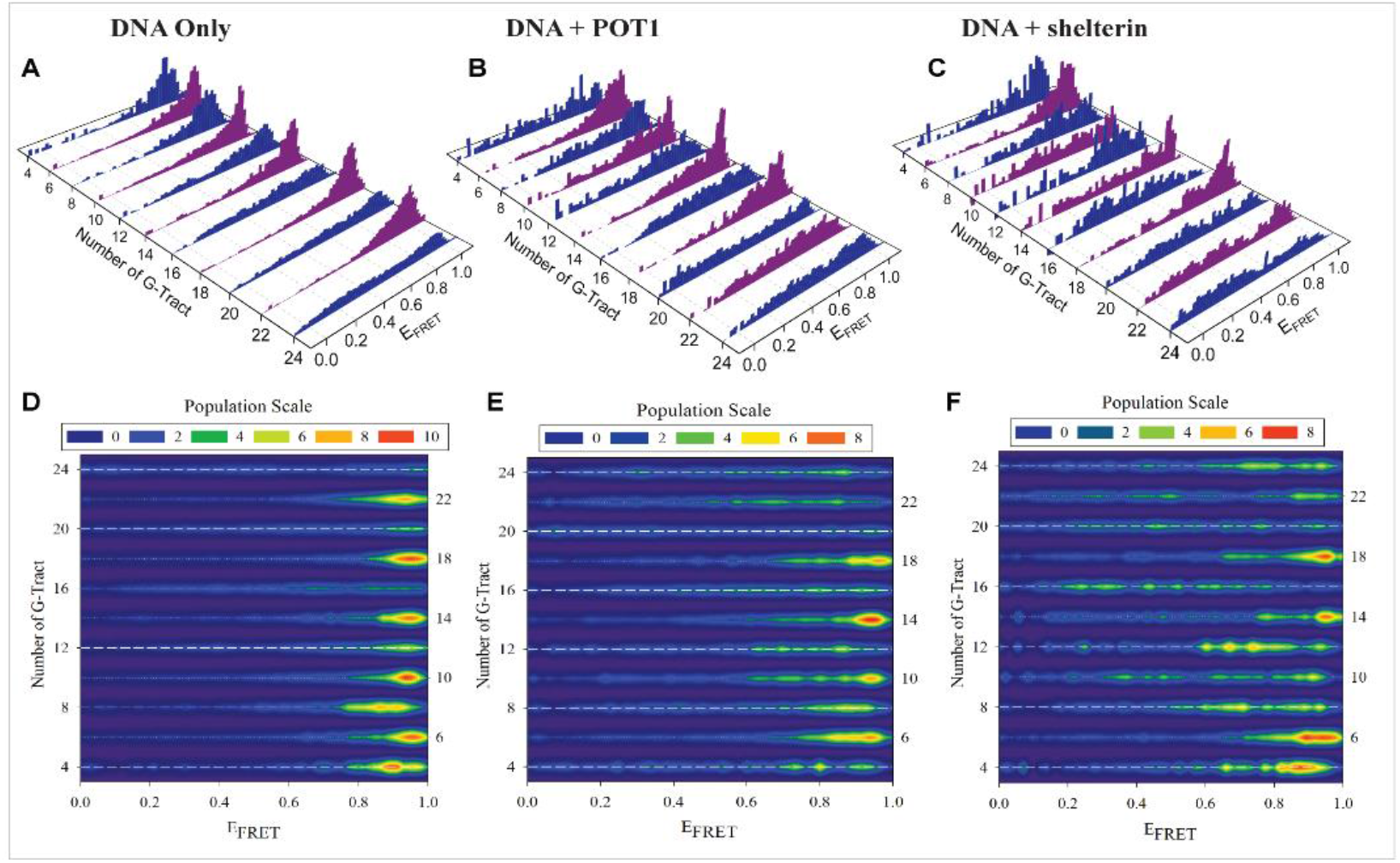
Normalized FRET histograms and contour plots for telomeric overhangs with 4-24 G-Tracts. FRET histograms are constructed from Cy5-PNA binding to different regions of ssTEL A) for DNA only, B) in the presence of POT1 and C) in the presence of shelterin (POT1, TPP1, TIN2 & TRF1). (D, E & F) Contour plots of histograms in A, B and C, respectively. Regions with concentrated red color are highly accessible while broad yellow/green regions indicate lower accessibility to a broader segment of ssTEL.

To quantify the broadness of FRET distributions, we introduce an S-parameter which is a measure of the uncertainty in determining where on the overhang the Cy5-PNA binds (Fig. 3). Broader distributions have a higher S-parameter since the binding sites are distributed over a more extensive part of the overhang, increasing the uncertainty in determining the binding location. In the absence of proteins, the S-parameter is maximum at [4n]G-Tract constructs and minimum at [4n+2]G-Tract constructs, with large variations between these two groups of constructs. These variations significantly diminish in the presence of POT1 and are almost eliminated by shelterin. These changes in the S-parameter distributions can be quantified by the variance of the S-parameter data, which shows a minimum in the presence of shelterin (σ^2^=6.5×10^−3^) and maximum in the absence of proteins (σ^2^=6.2×10^−2^).

**Figure 3:**
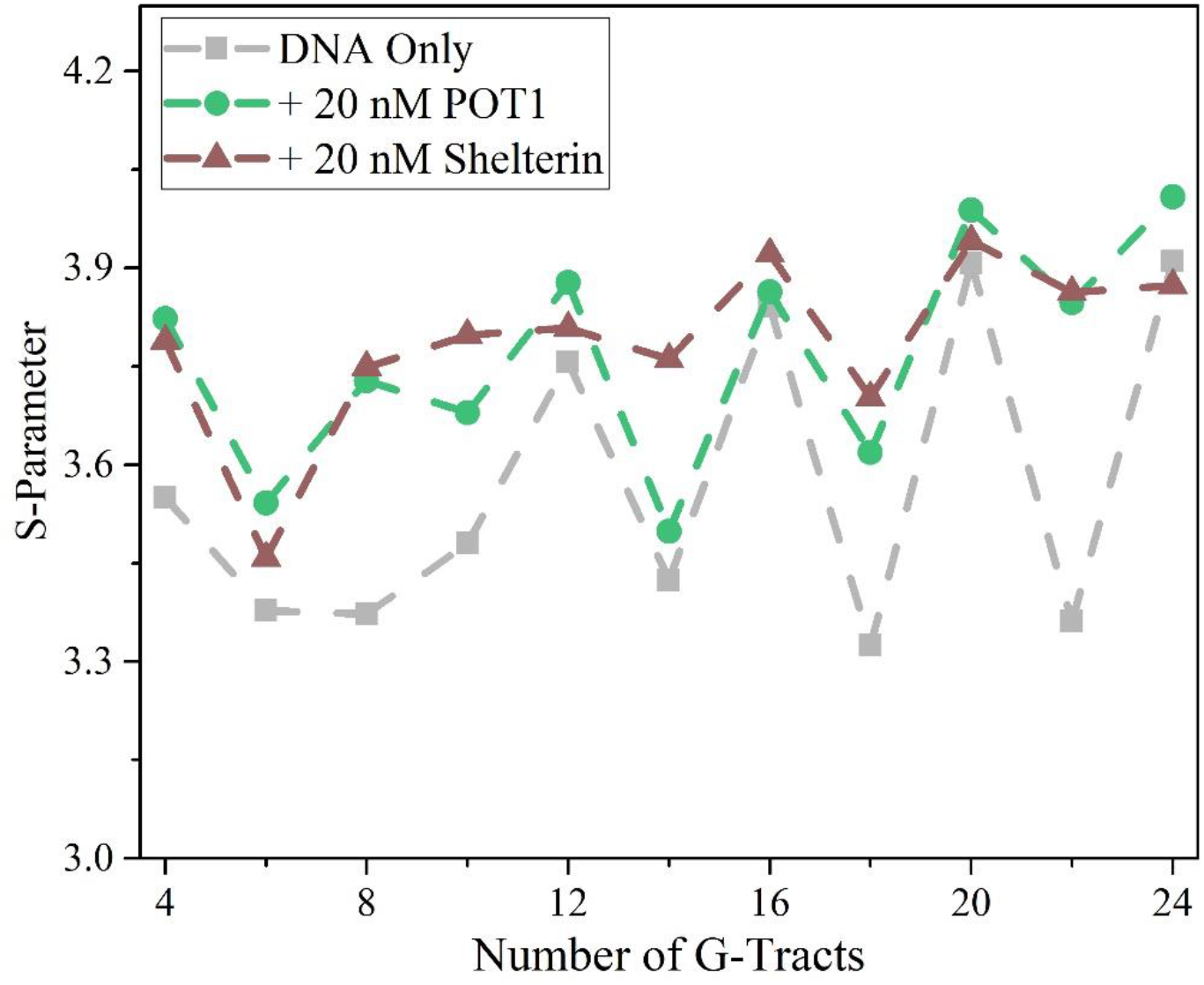
Scatter plot of S-parameter = -Σ_*i*_ *p*_*i*_ln(*p*_*i*_) in the absence of proteins (DNA-only) or presence of POT1 or Shelterin. S-parameter measures the broadness of FRET distributions. Larger S-parameters represent distribution of Cy5-PNA binding sites over a greater segment of ssTEL. The variation between the S-parameters for different constructs (different number of G-Tracts) decreases in the presence of POT1 and shelterin, i.e. S-parameters flatten, suggesting the accessibilities of different constructs become more uniform.

To quantify telomere accessibility, we calculated the Cy5-PNA binding frequency for different overhang lengths in the presence or absence of proteins (Fig. 4A). The binding frequencies significantly decrease (One-way ANOVA, p<0.001) in presence of POT1 or shelterin compared to their absence. This reduction was quantified by calculating the “relative accessibility”, i.e. the ratio of the binding frequencies observed in the presence of proteins to those observed in their absence (Fig. 4B). Mean relative accessibilities was reduced by ∼2.5 fold in the presence of 20 nM POT1 and ∼5 fold in the presence of 20 nM shelterin.

**Figure 4:**
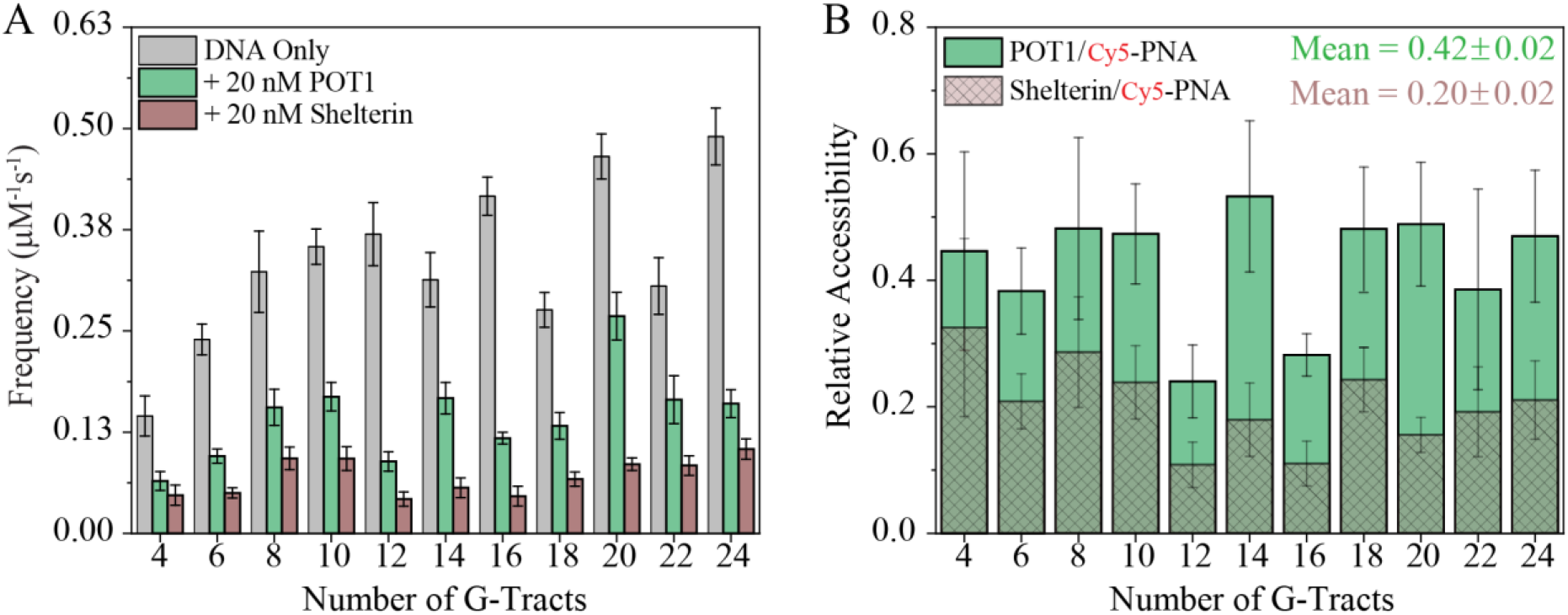
Cy5-PNA binding frequencies and relative accessibility of telomeric DNA overhangs. A) Binding of Cy5-PNA is quantified by analyzing the binding events in the absence of proteins and in the presence POT1 or shelterin. B) Relative accessibility represents the ratio of binding frequencies in the presence and absence of proteins. Mean accessibility is reduced ∼2.5 fold in the presence of POT1 and ∼5 fold in the presence of shelterin.

We also investigated whether the connection between POT1 (bound to ssTEL) and TRF1 (bound to the dsTEL) via TPP1 and TIN1 might impact the protection of the telomeric overhang. We performed measurements in the presence of POT1 and TRF1, without TPP1 and TIN2 which are required for establishing a connection between POT1 and TRF1 (Fig. 5A-C). These measurements demonstrate that TRF1 alone does not reduce accessibility of ssTEL (at the 0.05 level mean binding frequencies of TRF1 case are not significantly different from DNA-only case, One-way ANOVA). Also, adding TRF1 along with POT1 does not reduce the accessibility beyond that provided by the POT1-only case (at the 0.05 level mean binding frequencies of TRF1+POT1 case are not significantly different from POT1-only case, One-way ANOVA). These results suggest the connection between TRF1 and POT1, via TPP1 and TIN2, is critical in achieving the protection provided by the shelterin. Supplementary Fig. S4 shows the FRET histograms corresponding to these measurements.

**Figure 5:**
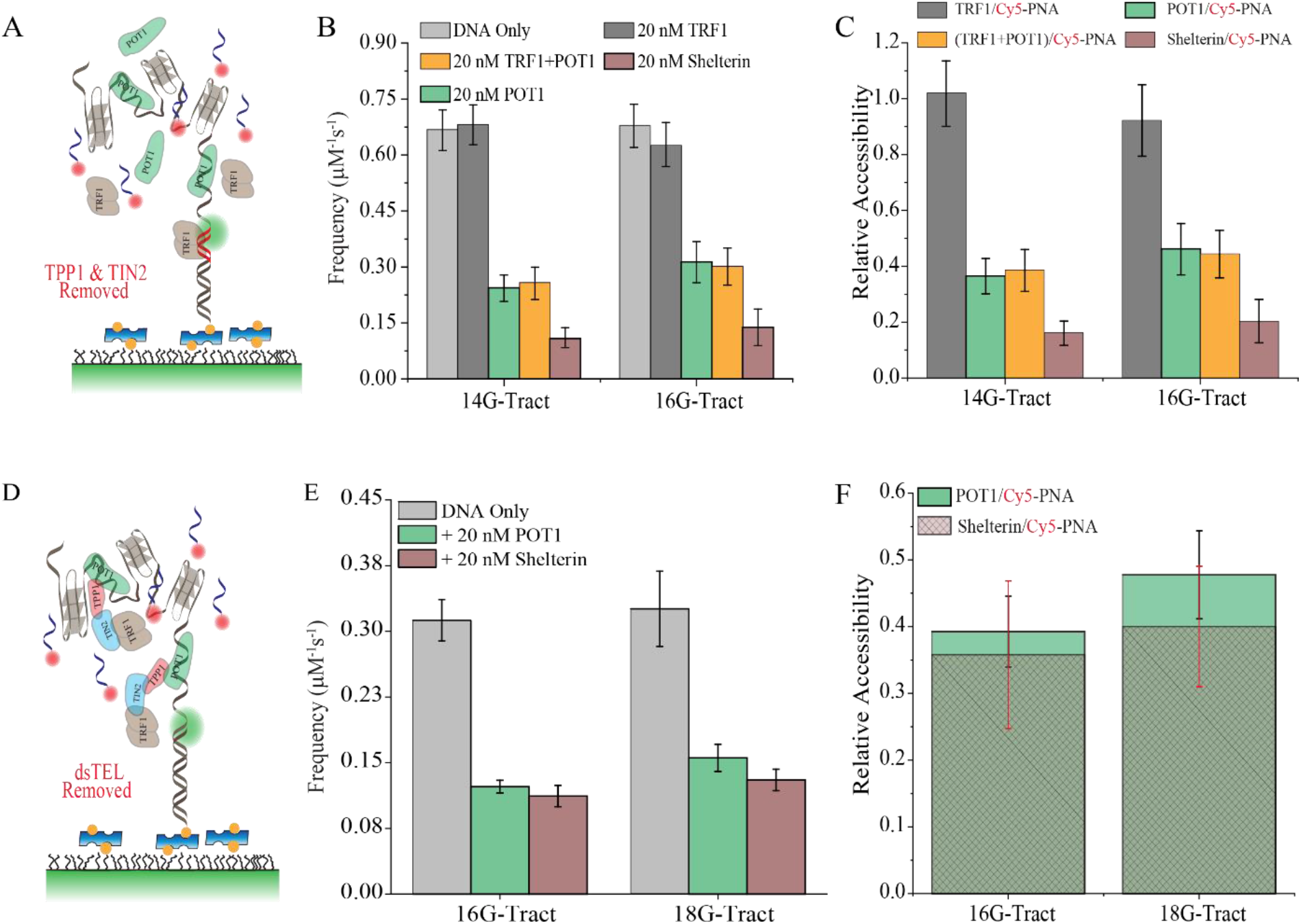
FRET-PAINT measurements probing the role of the connection between members of shelterin on ssTEL and dsTEL on overhang accessibility. (A) Schematics of FRET-PAINT assay where POT1 and TRF1 are used in the absence of TPP1 and TIN2. (B) Binding frequencies for 14G-Tract and 16G-Tract constructs. Frequencies for TRF1-only case are similar to those of DNA-only case while those of POT1 and TRF1 are similar to those of POT1-only case. (C) Relative accessibility for 14G-Trct and 16-G-Tract constructs. Relatively accessibility of the case when both TRF1 and POT1 are included is significantly less than that of shelterin. (D) Schematics of FRET-PAINT assay for modified DNA constructs that do not contain dsTEL (red duplex segment in Fig. 1 is removed). (E) Binding frequencies for modified 16G-Tract and 18G-Tract constructs (lacking dsTEL) in the absence of proteins and in the presence of POT1 or shelterin (POT1, TPP1, TIN2 and TRF1). F) Relative accessibility for modified 16G-Tract and 18G-Tract constructs. Relativity accessibility of shelterin is similar to that of POT1, suggesting the dsTEL is critical for the enhanced protection provided by shelterin.

To further explore this issue, we created analogs of 16G-Tract and 18G-Tract constructs that do not contain dsTEL (Fig. 5D), hence preventing binding of TRF1/TIN2 to this region. Fig. 5E shows the binding frequency of Cy5-PNA to these modified DNA constructs in the presence or absence of proteins. The relative accessibilities for these modified constructs in the presence of POT1 were similar to those for unmodified constructs (Fig. 5F). However, in the absence of dsTEL tracts, shelterin does not reduce the accessibility any more than that of the POT1-only case, i.e. ∼2.5 fold. These results demonstrate that the accessibility of telomere overhangs is reduced not only by POT1 but also by the connection between proteins that are bound to ssTEL and dsTEL.

We also tested if TRF2, instead of TRF1, in the shelterin would make a difference in binding frequencies or relative accessibilities of telomere overhangs. We repeated similar measurements with shelterin containing TRF2 (TRF2, TIN2, POT1, TPP1) using 14G-Tract and 16G-Tract constructs (Supplementary Fig. S5-S6). These measurements did not show a significant difference between TRF1 or TRF2 containing shelterin, further supporting our conclusion that connection between ssTEL and dsTEL is sufficient to reduce the accessibility of ssTEL overhang.

## DISCUSSION

Our comparative FRET-PAINT measurements on long telomeric overhangs provide important insights about how the distribution (location of accessible sites) and frequency (level of accessibility) of Cy5-PNA binding events are impacted by POT1 or shelterin. The observed accessibility patterns in the absence of proteins in Fig. 2 are consistent with earlier work which interpreted these patterns in terms of elevated levels of folding frustration in [4n]G-Tract constructs, which results from combinatorial effects associated with finite length of the overhang and the position of folded GQs on the overhang ^39,40^. Interestingly, introducing POT1 and shelterin largely eliminated the differences between different constructs (flattened the S-parameters). These observations suggest that POT1 binds more efficiently to the sites that are otherwise most accessible to Cy5-PNA and inhibits its binding to these sites. The further flattening of the S-parameter in the presence of shelterin indicates that members of shelterin that are localized on dsTEL also play a role in the protection of the overhang. Because other shelterin components do not directly bind ssTEL, it is not clear whether TRF1, TIN2, and TPP1 would have any impact on accessibility of ssTEL. However, we observed an additional 2-fold reduction in the presence of shelterin compared to POT1-only case, which we propose to be due to reduced accessibility of the region between dsTEL and ssTEL in the presence of shelterin. This region of the overhang was demonstrated to be the most accessible region of ssTEL ^39^ and a potential site for the connection between POT1 and TRF1. Therefore, binding of shelterin to this junction region might be an effective way to enable a robust protection of ssTEL. Interestingly, destabilization of GQs that might form in this region due to their proximity to dsTEL, which would have caused elevated levels of exposure for ssTEL, might be facilitating establishment of this critical connection between members of the shelterin. The measurements in Fig. 5 further support this proposal as they demonstrate that eliminating the dsTEL from DNA constructs or breaking the link between POT1 and TRF1 by eliminating TPP1 and TIN2 eliminates the enhanced protection provided by shelterin compared to POT1-only case.

The reduction in Cy5-PNA binding frequency (Fig. 4) in the presence of POT1 was not readily anticipated. It is known that POT1 has a mild GQ destabilization activity ^27,37,41,42^, which would have resulted in elevated exposure of ssTEL to Cy5-PNA unless such exposed sequences are effectively covered by POT1. Considering the 7-nt complementarity of Cy5-PNA and ssTEL and the minimum 9-nt binding site of POT1 (5′-TAGGGTTAG) ^11^, a tight competition between the two would have been expected. However, the ∼2.5 fold reduction in Cy5-PNA accessibility in the presence of POT1 suggests coverage of exposed telomeric sequences by POT1 is effective enough to exclude even this very short probe.

In this single molecule investigation, we demonstrated that a coordinated effort between members of the shelterin that bind to ssTEL and dsTEL reduces the accessibility of telomeric overhangs by ∼5-fold. To our knowledge this is the first demonstration of this protection mechanism on physiologically relevant overhang lengths. We also showed that the connection between POT1 and TRF1, via TPP1 and TIN2, is required for the shelterin induced protection of the overhang. These results suggest a consistent and compelling picture where the nature and strength of interactions between shelterin proteins and telomeric DNA complement the thermodynamic stabilities and folding characteristics of the tandem telomeric GQs. This complementarity, in return, enables effective protection of these critical regions of the genome.

## METHODS

### Preparation of DNA Constructs

Partial duplex DNA (pdDNA) constructs were formed by annealing a long strand that contains the telomeric overhang with a short strand (30 nt) that has a biotin at 3′-end and a Cy3 at 5′-end. Long DNA oligomers were purchased from Eurofins Genomics and were purified in-house using denaturing polyacrylamide gel electrophoresis (PAGE). The purification data is shown in Supplementary Fig. S1 and DNA sequences are given in Table S1. The short strand contains a TAACCCTAACCC sequence at its 5’-side which hybridizes with the complementary sequence on the long strand to create a 12 bp long dsTEL segment, which enables binding of TRF1 and connecting TIN2/TRF1 and POT1/TPP1. To illustrate, a pdDNA construct that contains 18 repeats of GGGTTA sequence at its overhang was formed by annealing a long strand <u>TGGCGACGGCAGCGAGGC</u> *TTAGGGTTAGGG* **TTA (GGGTTA)**_**18**_**G** with the short strand Cy3-*CCCTAACCCTAA* <u>GCCTCGCTGCCGTCGCCA</u>-Biotin. The italic sequences are complementary to each other and form the dsTEL segment that enables binding of TRF1 or TRF2. Similarly, the underlined sequences are complementary to each other and form an 18-bp duplex that separates the telomeric sequences from the surface. The bold sequences constitute the telomeric overhang (ssTEL) that has 18 repeats of GGGTTA sequence. For brevity, each GGGTTA segment will be referred to as a “G-Tract”. The pdDNA construct above will be called “18G-Tract” construct.

For annealing, the short strand and the long strand were mixed in a molar ratio of 1:4 in a buffer that contains 150 mM KCl and 10 mM MgCl_2_. This high MgCl_2_ concentration was used only during annealing assay, while all FRET-PAINT measurements were performed at 2 mM MgCl_2_. The annealing reaction was performed in a thermocycler by heating the samples at 95 °C for 3 minutes and then gradually cooling them to 30 °C in steps of 1 °C/3 min.

HPLC purified PNA strands labeled with Cy5 were purchased from PNA-Bio Inc. The Cy5-PNA employed in this study targets a 7 nt sequence, which includes a single G-Tract, on ssTEL. The sequence of PNA was <u>TAACCCT</u>T-Cy5, in which the underlined nucleotides are complementary to a telomeric sequence AGGGTTA. This probe is significantly shorter than PNA strands typically used for targeting and destabilizing GQs ^43^. The melting temperature (T_m_) of the duplex formed by Cy5-PNA and telomeric DNA (12.7±0.3 °C) is much lower than T_m_ = 69 °C ^44^ for a telomeric GQ under similar ionic conditions. Therefore, the Cy5-PNA does not introduce a significant destabilization to the telomeric GQs.

### Protein Constructs

POT1 and 4-component shelterin (POT1, TPP1, TIN2 and either TRF1 or TRF2) were expressed in insect cells as previously described ^45^. Briefly, human POT1 with an N terminal ZZ affinity tag, TEV cleavage site, and YBBR labeling site was cloned into an Omnibac vector, and the four-component shelterin containing either TRF1 or TRF2 was cloned into a BigBac vector, where POT1 was given an N terminal YBBR tag, POT1, TRF1 and TRF2 were each given an N terminal ZZ affinity tag and a TEV cleavage site, and TIN2 and TPP1 were each given an N-terminal His-MBP affinity tag and a TEV cleavage site. Protein was purified from insect cells as previously described ^46^. Briefly, plasmids containing genes of interest were transformed into DH10Bac competent cells (Berkeley MacroLab), and Bacmid DNA was purified using ZymoPURE miniprep buffers (Zymo Research, D4210) and ethanol precipitation. Insect cells were transfected using Fugene HD transfection reagent (Promega, E2311). The virus was amplified in progressively larger cultures. 1 mL of the P1 virus was used to infect 50 mL of Sf9 cells at 1 million cells/mL for 72 h. 10 mL of the P2 virus was used to infect 1 L of Sf9 cells at 1 million cells/mL and expression proceeded for 72 h. Cells expressing the protein of interest were harvested at 4,000 g for 10 min and resuspended in 50 mL lysis buffer (50 mM HEPES pH 7.4, 1 M NaCl, 10% glycerol, 1 mM PMSF, 1 mM DTT, and 1 tablet of protease inhibitor (Sigma, 4693132001)). Lysis was performed using 15 loose and 15 tight plunges of a Wheaton glass dounce. The lysate was clarified using a 45 min, 360,000 g spin in a Ti70 rotor. The supernatant was incubated with 1 mL IgG beads (IgG Sepharose 6 Fast Flow, GE Healthcare, 17096902) for POT1 or 1 mL amylose beads (New England BioLabs, E8021S) for shelterin for 1 hour. Beads were washed with 40 mL of lysis buffer followed by 40 mL of storage buffer (50 mM Tris pH 7.5, 300 mM KCl, 2 mM MgCl2, 10% glycerol, 1 mM DTT). Beads were then collected and incubated with TEV protease (Berkeley Macrolab, Addgene #8827) for 1 h at room temperature to elute the protein. Cleaved protein was run through a Superdex 200 Increase 10/300 GL size exclusion column (Cytiva, 28-9909-44) to remove TEV, cleaved affinity tags, and shelterin subcomplexes.

### Single-molecule FRET assay and measurements

A home-built prism**-**type total internal reflection fluorescence (TIRF) microscope built around an Olympus IX-71 inverted microscope was used for these measurements following the protocols described in reference ^47^. Laser drilled quartz slides were used after they were thoroughly washed with acetone and 1M potassium hydroxide (KOH), followed by piranha etching for 20 min. After surface functionalization with amino silane for 20-30 min, the slides and coverslips were initially passivated with a mixture of mPEG (PEG-5000, Laysan Bio, Inc.) and biotin PEG (biotin-PEG-5000, Laysan Bio, Inc.) in the molar ratio of 40:1 and kept overnight. The slides and coverslips were cleaned and dried with nitrogen gas and stored at -20 °C for future experiments. Before the experiments, the slide and coverslip were passivated with 333 Da PEG for 30-45 min to increase the density of the PEG brush. Finally, the microfluidic chamber was created between slide and coverslip by placing double-sided tape between them followed by sealing the chamber with epoxy. In addition, the microfluidic chamber was treated with 2% (v/v) Tween-20 to reduce non-specific binding. Following the removal of excess detergent from the chamber, 0.01 mg/mL streptavidin was incubated in the chamber for 2 minutes. Subsequently, the freshly annealed pdDNA samples diluted to 10-20 pM in 150 mM KCl and 2 mM MgCl_2_ were immobilized for 2-5 min on the surface which resulted in surface density of ∼300 molecules per imaging area (∼50 μm × 100 μm). Excess or unbound DNA was removed from the chamber by washing the chamber with a buffer that contains 150 mM KCl and 2 mM MgCl_2_. All FRET-PAINT measurements were carried out in an imaging buffer (50 mM Tris-HCl (pH 7.5), 2 mM Trolox, 0.8 mg/mL glucose, 0.1 mg/mL glucose oxidase, 0.1 mg/mL bovine serum albumin (BSA), 2 mM MgCl_2_, 150 mM KCl and 40 nM Cy5-PNA). In experiments with proteins, 20 nM POT1 or 20 nM shelterin were also included in the imaging buffer. The Cy5-PNA strand was heated to 85 °C for 10 minutes before it was added to the imaging buffer.

### FRET-PAINT measurements with proteins

The initial stock of POT1 (221 μM) and shelterin (POT1, TPP1, TIN2 and TRF1 at 5.5 μM) were aliquoted at a concentration of 2 μM and at a volume of 2 μl in a buffer that contains 50 mM Tris (pH 7.5), 300 mM KCl, 2 mM MgCl_2_ and 10% glycerol. The aliquots were flash frozen and stored at -80 °C for future use. Prior to each experiment the proteins were thawed in ice for 2-5 min and diluted to desired concentration in the imaging buffer (ingredients given above). The imaging buffer that contains proteins was incubated in the chamber for 10 min before data acquisition. After a set of FRET-PAINT measurements was completed, the chamber was incubated with 150 mM KCl and 100 mM MgCl_2_ for 15 min to ensure any remnant proteins dissociate from the DNA. The chamber was then washed with 1000 μL of 150 mM KCl and 2 mM MgCl_2_ to remove these remnant proteins. After waiting for 10 min for refolding of the GQs, new proteins were added to the chamber. As shown in Supplementary Fig. S3, this procedure enables removal of bound proteins and allows the telomeric overhang to attain the original folding pattern. To avoid variations between surface qualities in different microfluidic chambers to impact comparative studies, the measurements for a given pdDNA were first performed in the absence of proteins in a microfluidic chamber, followed by data acquisition in the presence of POT1 and then shelterin in the same chamber.

### Data acquisition and analysis

The donor fluorophore was excited with a green laser beam (532 nm) and the fluorescence signal was collected by an Olympus water objective (60x, 1.20 NA). An Andor Ixon EMCCD camera was used to record 1500-2000 frame long movies with 100 ms frame integration time. The recorded movies were processed and analyzed by a custom C++ script and finally a MATLAB code was used to generate single molecule time traces of donor and acceptor intensities. The single molecule time traces were further screened by a custom MATLAB script. For each selected single molecule time trace, the background was subtracted based on the remnant donor and acceptor intensities after donor photobleaches. The selected single molecule time traces were used to generate the FRET histograms. FRET efficiency was calculated by 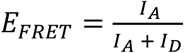 where *I*_*A*_ is the acceptor intensity and *I*_*D*_ is the donor intensity. Molecules that did not show any binding event contributed to the donor only (DO) peak at E_FRET_ = 0.06. The DO peak was subtracted from the FRET histograms which were then rescaled such that DO peak corresponds to E_FRET_ = 0.00. The number of molecules used in each FRET histogram is given in Table S2.

### Binding frequency analysis

An automated and bias-free step detection program, AutoStepfinder ^48^, was used to determine the E_FRET_ levels and dwell times in the single molecule time traces that were used to construct the FRET histograms. The Cy5-PNA binding frequencies were determined based on the Autostepfinder fits and were calculated by dividing the number of binding events to the total observation time. The total observation time is the sum of each individual time trace that show at least one binding event or donor photobleaching. The error bars for frequency analysis were determined by Bootstrapping analysis ^49^ with a sample size of 2000 and 95% confidence level ^32^.

### S-parameter calculation

The population of each of the FRET histograms was normalized to 100%. The broadness of FRET distributions was quantified by an S-parameter (Shannon entropy divided by Boltzmann constant): *S* = − Σ_*i*_ *p*_*i*_ ln *p*_*i*_, where *p*_*i*_ is the probability of a particular FRET level *i*. The *p*_*i*_ is obtained by dividing population at the corresponding FRET level by 100 (Fig. 4) so that 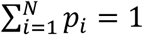 for a given FRET histogram.

## Acknowledgements

This work was supported by NIH (1R15GM123443 and R15GM146180 to H.B.). We thank Dr. Soumitra Basu for use of his lab to perform the PAGE experiments.

## Contributions

S.S., A.Y. and H.B. designed the study; S.S. and A.J. led the experimental work; A.J. prepared the protein constructs, G.M., S.K., M.E.H., and P.G. contributed to experimental work; S.S. led the data analysis, S.S., A. Y. and H.B. wrote the manuscript with input from all authors.

## Corresponding author

Correspondence to Ahmet Yildiz or Hamza Balci

## Ethics declarations Competing interests

The authors declare no competing interests.

