## Supplementary Information for "Impact of Shelterin Complex on Telomere Accessibility"

**DNA Sequences**

The sequence of the Cy5-PNA probe was TAACCCTT-Cy5, the underlined nucleotides being complementary to the telomeric sequence. The partial duplex DNA (pdDNA) constructs were created by annealing a 30-nt long stem strand with a longer strand that includes a complementary sequence to the stem strand and the telomeric overhang. The following strands were used to create the pdDNA constructs.

**Table S1:** Sequences used for creating pdDNA constructs. The nucleotides in green fonts constitute the overhang.

| Pd-DNA Constructs | Long Strand (Sequence in 5′–3′) |
|---|---|
| 4G-Tract | TGGCGACGGCAGCGAGGCTTAGGGTTAGGGTTA $(GGGTTA)_4$ G |
| 6G-Tract | TGGCGACGGCAGCGAGGCTTAGGGTTAGGGTTA $(GGGTTA)_6$ G |
| 8G-Tract | TGGCGACGGCAGCGAGGCTTAGGGTTAGGGTTA $(GGGTTA)_8$ G |
| 10G-Tract | TGGCGACGGCAGCGAGGCTTAGGGTTAGGGTTA $(GGGTTA)_{10}$ G |
| 12G-Tract | TGGCGACGGCAGCGAGGCTTAGGGTTAGGGTTA $(GGGTTA)_{12}$ G |
| 14G-Tract | TGGCGACGGCAGCGAGGCTTAGGGTTAGGGTTA $(GGGTTA)_{14}$ G |
| 16G-Tract | TGGCGACGGCAGCGAGGCTTAGGGTTAGGGTTA $(GGGTTA)_{16}$ G |
| 18G-Tract | TGGCGACGGCAGCGAGGCTTAGGGTTAGGGTTA $(GGGTTA)_{18}$ G |
| 20G-Tract | TGGCGACGGCAGCGAGGCTTAGGGTTAGGGTTA $(GGGTTA)_{20}$ G |
| 22G-Tract | TGGCGACGGCAGCGAGGCTTAGGGTTAGGGTTA $(GGGTTA)_{22}$ G |
| 24G-Tract | TGGCGACGGCAGCGAGGCTTAGGGTTAGGGTTA $(GGGTTA)_{24}$ G |
| Stem-30 | Cy3-CCCTAACCCTAA GCCTCGCTGCCGTCGCCA-biotin |
| Cy5-PNA | TAACCCTT-Cy5 |

The subscripts designate the number of repeats, i.e. $(GGGTTA)_4$ refers to GGGTTAGGGTTAGGGTTAGGGTTA. Stem-30 strand (30 nt) hybridizes with a segment of the long strand (30 nt on 5'-side) and creates a duplex DNA which is attached to the surface via biotin-streptavidin conjugation. The purple nucleotides of the long strands hybridize with the purple nucleotides of the Stem-30 strand and form a 12-bp long telomeric duplex region.

**Table S2**. Data statistics used in Fig 2 and 3.

| pdDNA Constructs | Conditions | Total number of molecules that passed single molecule screening | Number of molecules that show at least one binding event | Number of binding events |
|---|---|---|---|---|
| 4G-Tract | DNA-Only | 434 | 54 | 110 |
| | POT1 | 567 | 46 | 63 |
| | Shelterin | 576 | 26 | 50 |
| 6G-Tract | DNA-Only | 352 | 143 | 312 |
| | POT1 | 593 | 116 | 183 |
| | Shelterin | 523 | 49 | 74 |
| 8G-Tract | DNA-Only | 252 | 72 | 194 |
| | POT1 | 286 | 54 | 105 |
| | Shelterin | 468 | 52 | 93 |
| 10G-Tract | DNA-Only | 517 | 238 | 516 |
| | POT1 | 468 | 106 | 187 |
| | Shelterin | 380 | 56 | 63 |
| 12-Tract | DNA-Only | 492 | 149 | 522 |
| | POT1 | 416 | 73 | 122 |
| | Shelterin | 452 | 33 | 50 |
| 14G-Tract | DNA-Only | 339 | 116 | 278 |
| | POT1 | 459 | 94 | 187 |
| | Shelterin | 494 | 32 | 46 |
| 16G-Tract | DNA-Only | 684 | 325 | 1000 |
| | POT1 | 856 | 199 | 317 |
| | Shelterin | 801 | 68 | 76 |
| 18G-Tract | DNA-Only | 348 | 162 | 340 |
| | POT1 | 391 | 74 | 123 |
| | Shelterin | 540 | 66 | 92 |
| 20G-Tract | DNA-Only | 482 | 301 | 972 |
| | POT1 | 399 | 153 | 352 |
| | Shelterin | 535 | 81 | 109 |
| 22G-Tract | DNA-Only | 304 | 148 | 222 |
| | POT1 | 281 | 62 | 146 |
| | Shelterin | 373 | 57 | 78 |
| 24G-Tract | DNA-Only | 380 | 139 | 266 |
| | POT1 | 373 | 84 | 129 |
| | Shelterin | 431 | 40 | 53 |

**Gel Purification Data**

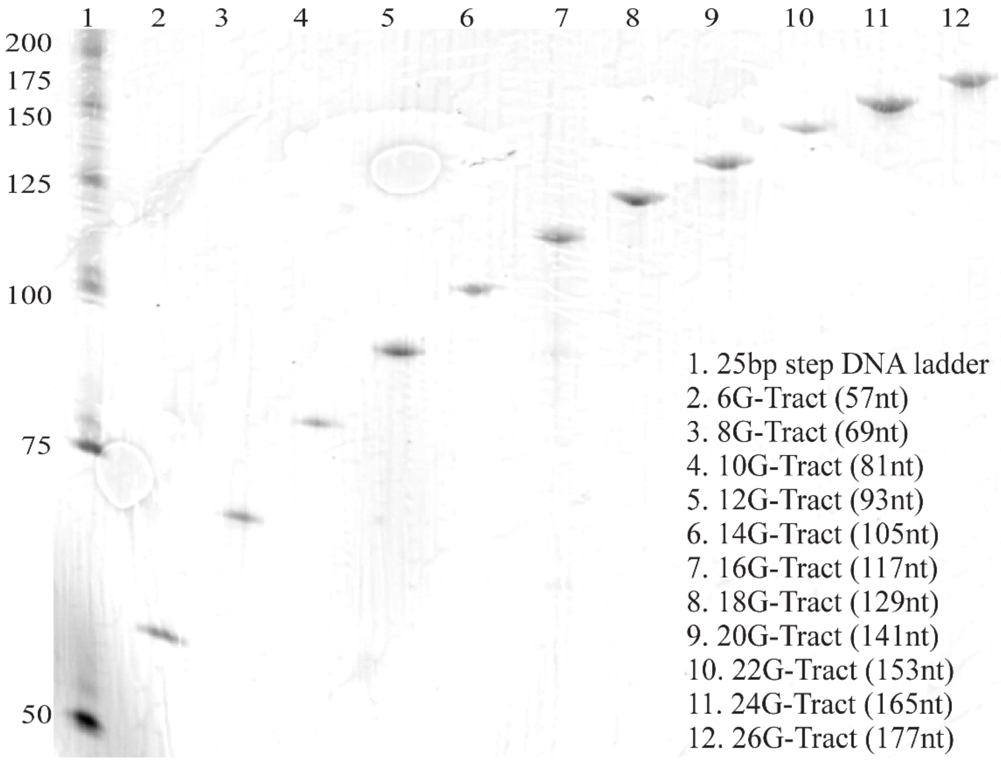


**Figure S1:** Denaturing polyacrylamide gel of purified DNA oligonucleotides. DNA oligos were purified via denaturing 10% polyacrylamide gel electrophoresis (PAGE). Full-length products were visualized by UV shadowing and the bands were excised from the gel. The DNA oligos were gathered via the crush and soak method by incubating and rocking of the gel slice overnight at room temperature in a solution of 300 mM NaCl, 10 mM Tris-HCl, and 0.1 mM EDTA (pH 7.4). The eluants were concentrated with sec-butanol and precipitated with 3 volumes of 100% ethanol. Salt was removed by washing twice, with 70% ethanol. After vacuum drying, the DNA pellet was dissolved in nuclease free water. Purified DNA oligos were resolved via denaturing 10% PAGE and the gel was stained with SYBR gold for 15 minutes. The gel was then visualized on a Typhoon FLA 9500 fluorescence imager (GE Life Sciences) by selecting Cy3 scanning mode.

### Extension of the Overhang by POT1

In order to quantify the stretching of the overhang by POT1 and ensure that detection sensitivity for Cy5-PNA binding is maintained, we designed a DNA construct that has a telomeric overhang which terminates with an 11-nt long non-telomeric sequence. We also designed a Cy5-labeled short imager strand that can bind only to an 8-nt segment of this non-telomeric sequence, ensuring we will measure the minimum possible FRET efficiency in the system as the imager strand can bind only to the 3'-end which is the furthest point from donor fluorophore (located at the ssDNA/dsDNA junction). Figure S2 shows these data where we performed FRET-PAINT measurements before and after introducing POT1 to a 24G-Tract construct. As expected, introducing POT1 shifts the FRET efficiency measured between the ends to lower values; however, the resulting histogram is still within the measurable FRET range. The overall compactness of the overhang, the scarcity of available binding sites for POT1, and low GQ unfolding activity of POT1 result in a relatively minor stretching of the overhang. The following DNA constructs were used in this study:

**24G-Tract-11nt:** TGGCGACGGCAGCGAGGCTTA GGGTTA GGGTTA GGGTTA GGGTTA GGGTTA GGGTTA GGGTTA GGGTTA GGGTTA GGGTTA GGGTTA GGGTTA GGGTTA GGGTTA GGGTTA GGGTTA GGGTTA GGGTTAG GGTTA GGGTTA GGGTTA GGGTTA GGGTTA GGGTTA G TACGATCGCAG
**Stem Strand**: Cy3-GCCTCGCTGCCGTCGCCA-biotin
**8nt DNA Imager Strand**: 5'-AGCGATCGT-Cy5-3'

Nucleotides with red fonts in pd-24G-Tract-11nt strand and in Stem Strand are complementary to each other and form an 18-bp long duplex. The green nucleotides of 8nt DNA Imager Strand are complementary to green nucleotides at the 3'-end of pd-24G-Tract-11nt construct. Transient bindings of this DNA imager strand result in FRET signals that are compiled in Figure S2-B.

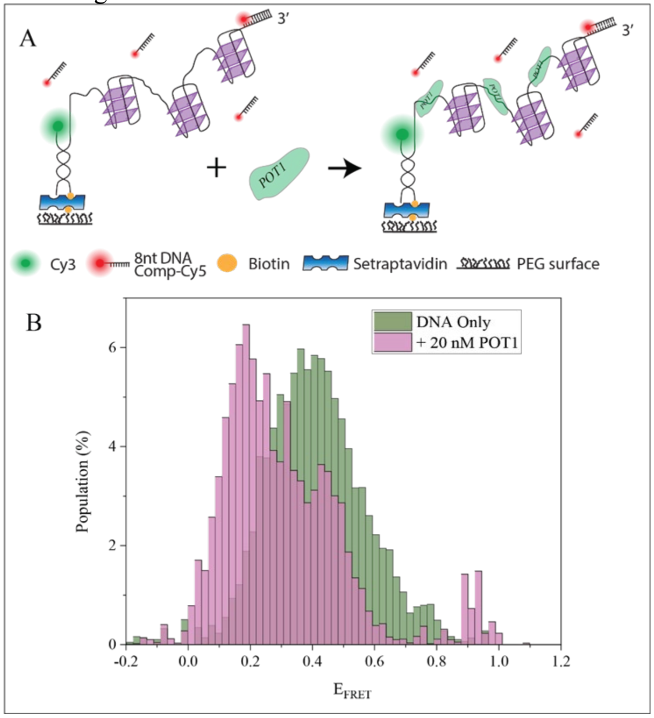


**Figure S2:** Stretching of telomeric overhang by POT1. A) The telomeric overhang used in this study (24G-Tract-11nt) has an additional non-telomeric 11 nt at the 3' end. The Cy5-labeled imager is complementary to 8 nt of this non-telomeric sequence. Binding of the imager strand to this non-telomeric region at the 3'-end results in a FRET signal that represents the lowest possible FRET value in FRET-PAINT assay. Binding of POT1 is expected to stretch the overhang and reduce the measured FRET efficiency. B) FRET histograms are generated by compiling these binding events to the 3' end in presence (brown) or absence (green) of POT1. The POT1 bound peak is 0.2, showing that even for the longest overhang used in this study (24G-Tract), the FRET is still in detectable range.

### Removal of POT1 and Recovering Folding Pattern of the Overhang

To avoid variations due to surface quality impact the comparative studies between the measurements performed in the absence of proteins or presence of POT1 or the Shelterin complex, all measurements for a given DNA construct were performed within the same sample chamber. We performed the measurements in the absence of proteins first, followed by POT1 measurements and then Shelterin measurements. After POT1 measurements, the chamber was incubated with a buffer that contains 150 mM KCl and 100 mM $MgCl_2$ for 10 min. The higher $MgCl_2$ concentration in this buffer helps with removing the POT1 from the overhang. The chamber was then washed with an imaging buffer that contains 150 mM KCl and 2 mM $MgCl_2$ and incubated in this buffer for another 10 min. At the end of this process, we checked whether the original folding patterns (those in the absence of proteins) were recovered. To directly probe the folding pattern, the donor-acceptor fluorophores were placed across the overhang, as shown in Figure S3A-B. Figure S3C shows these data for an overhang that contains 3G-Tracts and Figure S3D shows the data for overhang with 10G-Tracts.

The following DNA sequences were used in these measurements:
pd-3G-Tract: 5'-TGGCGACGGCAGCGAGGCTATTATTAGGGTTAGGGTTAGGGTTAG-Cy3-3'
pd-10G-Tract: TGGCGACGGCAGCGAGGCTTA $(GGGTTA)_{10}$G-Cy3-3'
Stem strand: Cy5-GCCTCGCTGCCGTCGCCA-biotin

Nucleotides with red fonts of pd-3G-Tract and pd-10G-Tract sequences are complementary to the nucleotides of the stem strand in blue fonts and form duplex part of the pdDNA constructs after annealing.

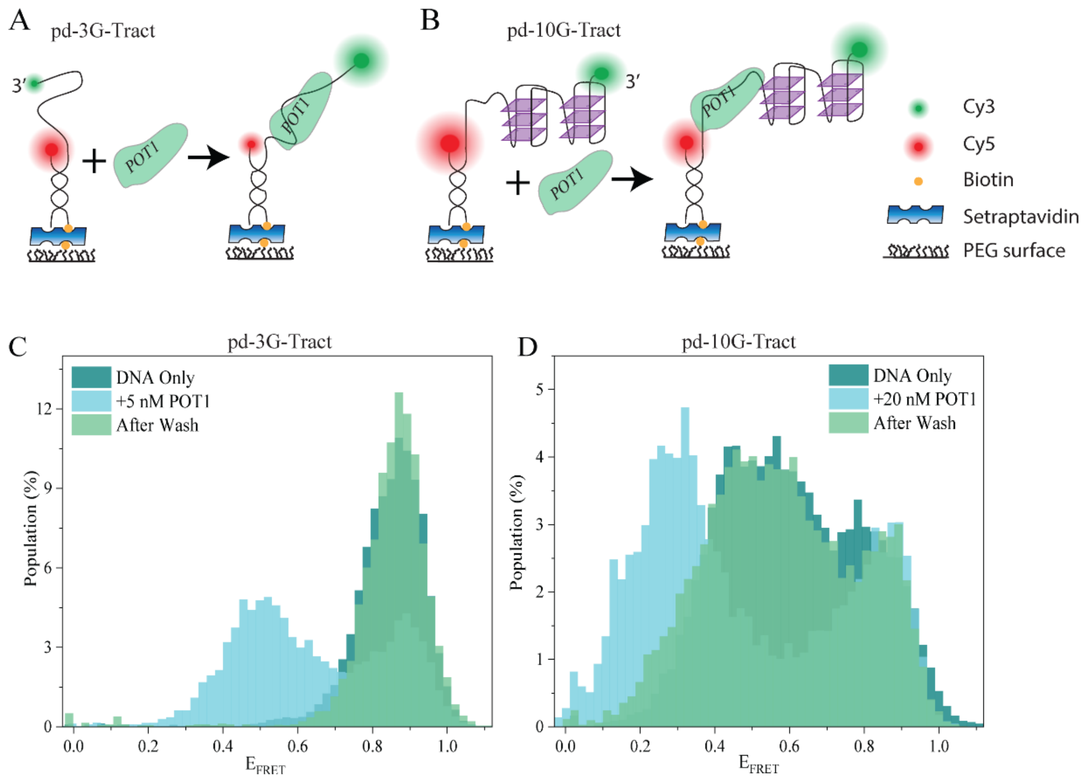


**Figure S3:** Schematics of smFRET assay for dual (Cy3/Cy5) labelled constructs of the (A) 3G-Tract construct and (B) 10G-Tract construct. These constructs are designed for traditional smFRET measurements (rather than FRET-PAINT measurements where the acceptor was placed on the Cy5-PNA strand) to directly probe the folding patterns. Two pdDNA constructs with overhang length of 3G-Tract (A) or 10G-Tract (B) are immobilized on the surface. The donor fluorophore is attached on the long strand at the 3′ end and acceptor fluorophore is at the 5'-end of the short strand. C) and D) Normalized FRET histograms for pdDNA constructs with 3G-Tract overhang or 10G-Tract overhang in the presence or absence of POT1. Adding POT1 results in a low FRET peak in both 3G-Tract and 10G-Tract constructs. After following the protocol to wash out the proteins, the original FRET pattern in recovered. Therefore, we conclude that the protocol we employed enables removal of the bound POT1 and refolding of the GQ.

### Measurements with POT1 and TRF1

In order to test whether the connection between POT1 and TRF1, created by TIN2 and TPP1, would impact the accessibility of telomeric overhang we performed measurements with POT1 and TRF1, while TIN2 and TPP1 were excluded from these measurements. The binding frequencies and relative accessibilities for these measurements are given in Figure 5A-C. Figure S4 shows the relevant accessibility patterns for these measurements. Similar to the observations in terms of binding frequencies, the accessibility pattern for TRF1-only case is very similar to that of DNA-only case. So TRF1 alone does not impact where the Cy5-PNA probe binds and how frequently it binds. In addition, the accessibility pattern in the presence of TRF1 and POT1 is very similar to that of POT1-only case, suggesting TRF1 does not impact the protection provided by POT1 in the absence of TIN2 and TPP1.

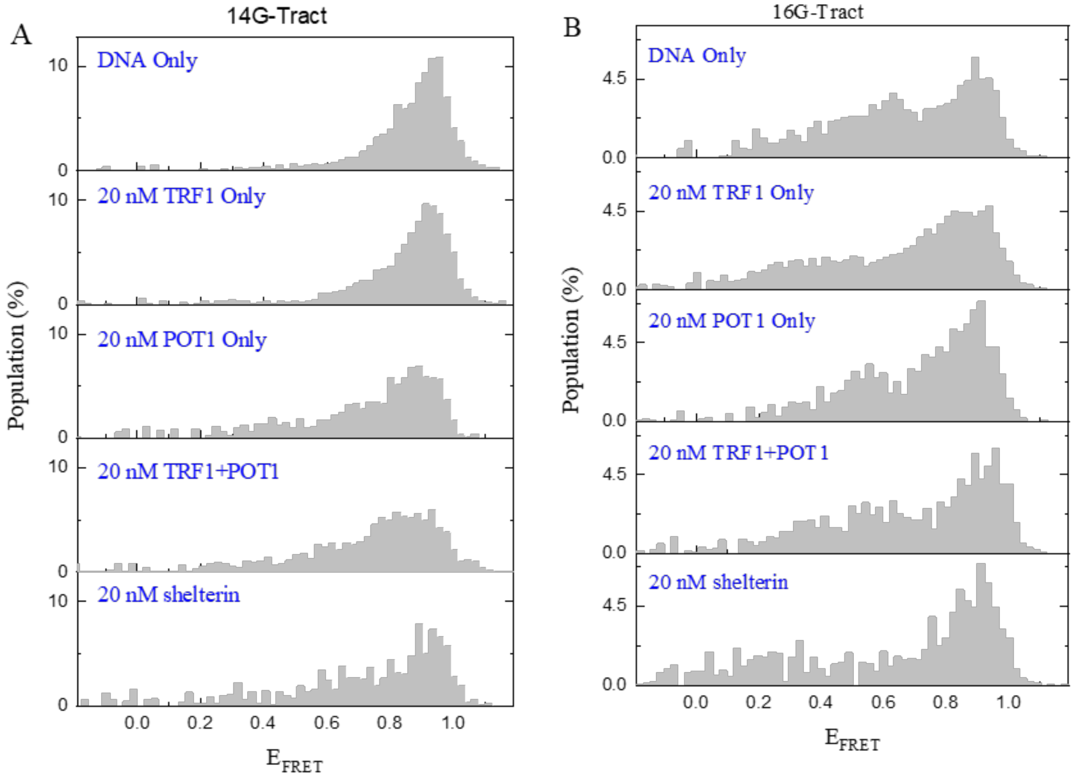


**Figure S4:** Accessibility patterns on 14G-Tract and 16G-Tract constructs. Excluding TIN2 and TPP1 breaks the connection between TRF1 and POT1 and prevents TRF1 to have any impact on the protection of telomeric overhangs.

**Measurements with Shelterin that Contains TRF2**

We repeated the FRET-PAINT measurements with a Shelterin complex that contained TRF2 instead of TRF1, i.e., POT1, TPP1, TIN2, and TRF2. The 14G-Tract and 16G-Tract constructs were used for these measurements which are shown in Figure S5-S6. Both the accessibility patterns (Figure S5) and the binding frequencies (Figure S6) are similar to those obtained for the shelterin that contains TRF1.

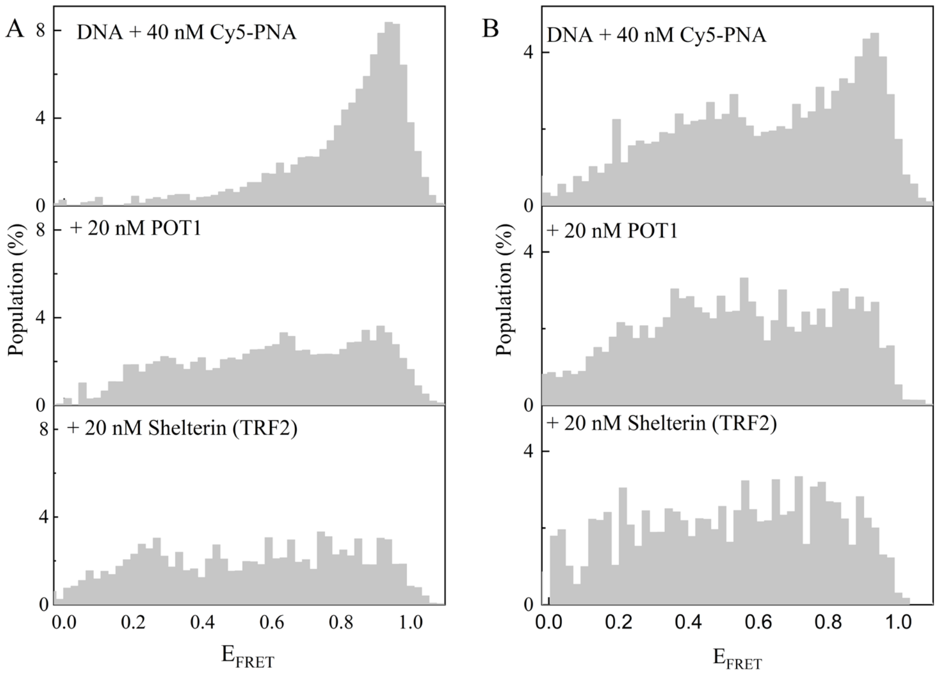


**Figure S5:** Normalized FRET-PAINT histograms showing accessibility patterns for (A) 14G-Tract and (B)16G-Tract constructs in presence and absence of proteins. The shelterin contained TRF2 instead TRF1 (POT1, TPP1, TIN2 and TRF2). These accessibility patterns are very similar to those obtained for the shelterin that contained TRF1.

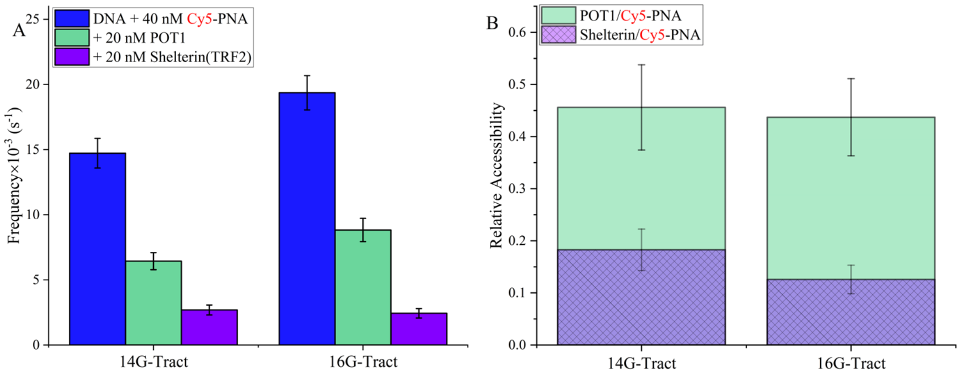


**Figure S6:** (A) PNA binding frequencies and (B) relative accessibilities for 14G-Tract and 16G-Tract constructs when a shelterin that contains TRF2, instead of TRF1, was used in the measurements. Relative accessibility represents the ratio of binding frequency in the presence versus the absence of proteins. Similar to the case of shelterin with TRF1, binding frequency is reduced by approximately 5-fold in the presence of shelterin that contains TRF2.
